# Aclarubicin disrupts RNA polymerase II progression at replication-coupled histone genes

**DOI:** 10.64898/2026.07.08.737292

**Authors:** Kevin K. Nguyen, Matthew Wooten, Kami Ahmad, Steven Henikoff

## Abstract

Anthracyclines are effective chemotherapeutic agents that cause DNA and chromatin damage. One member of the anthracyclines, aclarubicin, has gained therapeutic interest due to its ability to kill cancer cells through chromatin-based mechanisms, thus avoiding the off-target effects associated with DNA damage. However, the molecular mechanism of action leading to aclarubicin-induced chromatin damage remains elusive. Here we performed Cleavage Under Targets and Tagmentation (CUT&Tag) of RNA polymerase II (Pol II) and other transcriptional regulators in human cells following aclarubicin treatment. We found that aclarubicin strongly disrupts the replication-coupled histone genes, resulting in a nonproductive accumulation of Pol II transcription machinery here beyond the levels at other genes. We attribute this sensitivity to the dense Pol II loading and rapid transcription of the histone genes, which intensify the chromatin-disrupting effects of aclarubicin. Together, our findings shed light on the mechanism of aclarubicin as an anticancer drug and point to the replication-coupled histone genes as a target for therapeutic intervention.

## Introduction

Eukaryotic gene expression by RNA polymerase II (Pol II) occurs in the context of chromatin, the complex of DNA and histone proteins that forms the structural basis of chromosomes. The fundamental repeating unit of chromatin, the nucleosome, is composed of a segment of DNA wrapped around a histone octamer of one H3-H4 tetramer and two H2A-H2B dimers, which are incorporated into chromatin in a replication-coupled manner (*1–3*). On the surface of the nucleosome core particle, the linker histone H1 binds the sites where DNA enters and exits. Chromatin structure and Pol II transcription are tightly coordinated. Transcription elongation through gene bodies requires Pol II to associate with a host of transcription factors: some promote its processivity, while others displace nucleosomes ahead of the polymerase and reassemble them behind the enzyme, providing access to genetic information while preserving chromatin integrity. Beyond transcription, maintaining chromatin integrity is critical for cell proliferation. Every cell cycle, chromatin is duplicated and newly synthesized DNA must be assembled into nucleosomes. Cell proliferation therefore depends on the coupling of DNA replication and histone synthesis, where dysregulation leads to diseases such as cancer.

Treatments that target fundamental aspects of cell proliferation remain the basis of cancer therapy. While many chemotherapeutics target features of DNA replication, the anthracyclines have begun to highlight the broader potential of targeting chromatin duplication and structure. Anthracyclines are widely prescribed anticancer drugs that consist of a tetracyclic ring attached to an amino sugar and intercalate in between DNA base pairs at GC-rich regions. Compared with other intercalating molecules, anthracyclines have been proposed to have a broad range of mechanisms that span DNA damage, topoisomerase inhibition, and disruption of the transcriptional and chromatin landscape (*4, 5*). One member of the anthracyclines, aclarubicin, produces no detectable DNA damage while maintaining the same levels of potency as other anthracyclines such as doxorubicin (*6, 7*). Previous studies suggest that the efficacy of aclarubicin treatment may rest in its ability to induce chromatin damage, where its intercalation alters the biophysical properties of DNA and disrupts DNA–histone interactions (*8–10*). Nonetheless, the molecular consequences of aclarubicin-mediated chromatin damage in human cells—in particular, which genomic sites are most sensitive and how transcription is dysregulated at those sites—remain unclear.

Our prior work showed that aclarubicin increases the elongating form of Pol II in *Drosophila* cells (*10*). Therefore, we sought to understand whether these effects are conserved in human cells. In this study, we defined the effects of aclarubicin treatment on transcriptional activity in human K562 cells. Focusing on the effects of aclarubicin at the replication-coupled histone genes (hereafter referred to as histone genes), we found that aclarubicin redistributes Pol II into gene bodies and substantially accumulates elongating Pol II and the histone gene transcription factor NPAT at these sites. We showed that the aclarubicin-induced enrichment of transcription machinery at the histone genes is attenuated when we inhibit transcription initiation and is ultimately nonproductive. Collectively, this work points to the histone genes as sites highly sensitive to aclarubicin-induced transcriptional disruption.

## Results

### Aclarubicin treatment increases Pol II-Ser2p at active genes

To determine the impact of aclarubicin on transcriptional activity in human cells, we treated proliferating cultures of K562 cells and then measured cell numbers, viability, and size after 30 minutes, 4 hours, and 24 hours (fig. S1, A to C). We found substantial changes in cell proliferation after 24 hours while the cells maintained viability. Seeking to observe acute drug effects following aclarubicin treatment, we moved forward with a 30-minute treatment window as our main experimental timepoint (*10*).

The transition of Pol II from an initiating state to elongating is accompanied by phosphorylation changes on the Carboxy-Terminal Domain (CTD) of its largest subunit Rpb1. The Pol II CTD features 52 heptapeptide repeats (YSPTSPS) in mammals. Serine 5 phosphorylation (Ser5p) predominates at promoters while Pol II is paused and is maintained during the initial stages of transcription (*11, 12*). Serine 2 phosphorylation (Ser2p) accumulates as Pol II moves into gene bodies. To map these CTD modifications, we performed Cleavage Under Targets and Tagmentation (CUT&Tag) (*13*) targeting total Pol II, the Ser5p-marked form, and the Ser2p-marked form of Pol II at the transcription start sites (TSSs) of 19,305 annotated gene promoters in the hg19 human genome assembly. Total Pol II signal was enriched at the TSS of 8,272 promoters in control samples (Fig. 1, A and D). Total Pol II signal was also apparent at 1,036,067 of the 2,343,081 ENCODE-annotated human Candidate *cis*-Regulatory Elements (cCREs) in the human genome (*14*) (fig. S1D). Cells treated with aclarubicin for 30 minutes showed minimal changes in total Pol II binding at promoters or cCREs compared with controls (fig. S1E). The largest differences in total Pol II signal, both gain and loss, were confined to a minority of highly occupied regions.

**Fig. 1.**
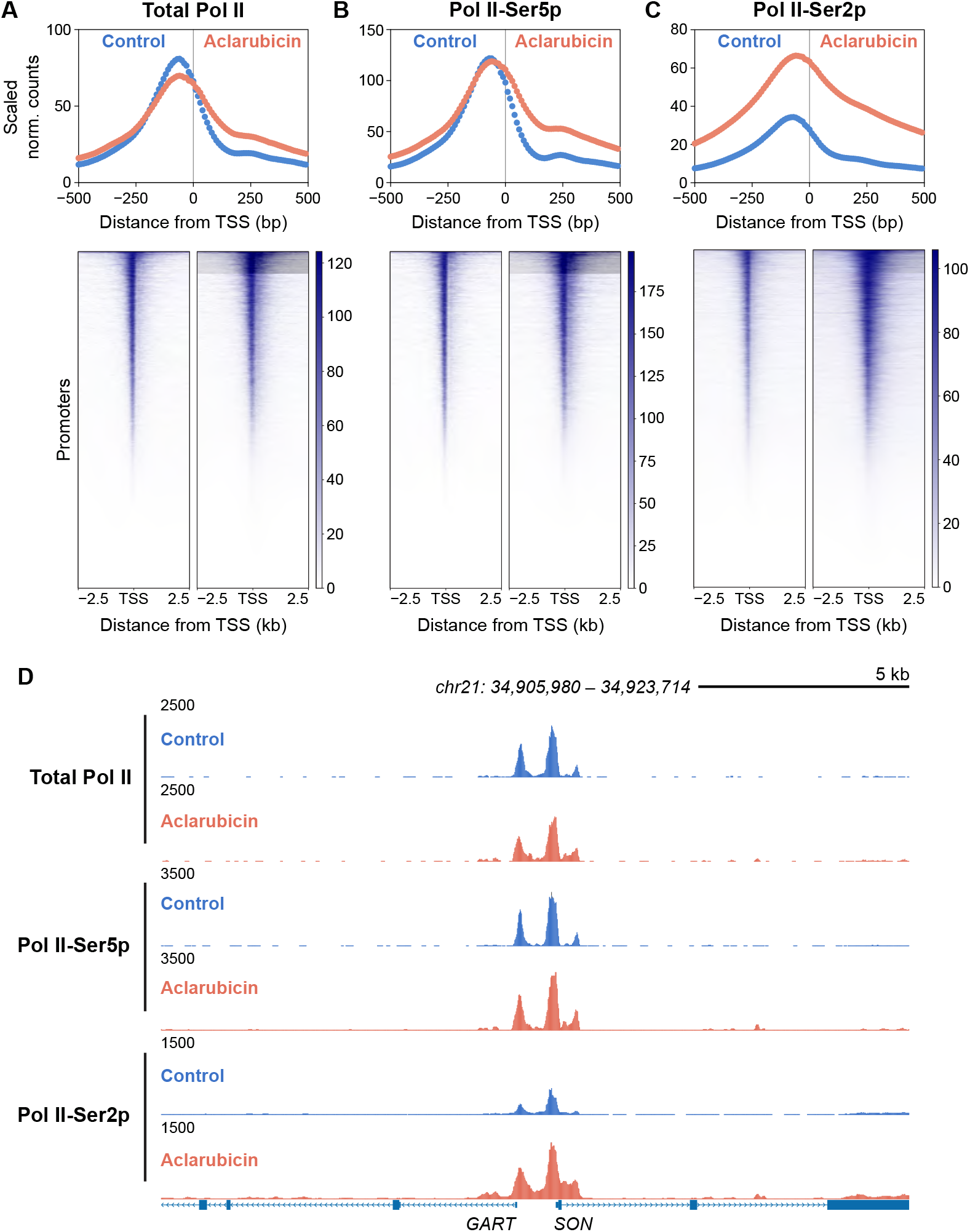
Aclarubicin treatment causes gains in Pol II-Ser2p. **(A to C)** Average signal distribution plot and heatmap of total Pol II (Rpb1) (A), paused Pol II (Ser5p) (B), and elongating Pol II (Ser2p) (C) CUT&Tag signal under dimethyl sulfoxide (DMSO) control and aclarubicin treatment conditions aligned to the transcription start site (TSS) of 19,305 hg19 promoters. For (A) and (B), merged data from two biological replicates per condition (control and aclarubicin-treated) shown. For (C), merged data from eight biological replicates per condition shown. **(D)** Representative genome browser distributions of Pol II under control and aclarubicin treatment conditions at *GART* and *SON* gene promoters.

For Pol II-Ser5p, we observed occupancy around the TSS of 9,282 promoters in control samples (Fig. 1, B and D). Pol II-Ser5p signal at TSSs was similar between control and aclarubicin-treated conditions. Analyzing the difference in Pol II-Ser5p signal between aclarubicin-treated and control conditions, we found minimal net change across cCREs and genes (fig. S1, F and G). This analysis also showed that the largest changes in Pol II-Ser5p signal were limited to regions of high Pol II-Ser5p occupancy.

Together, these data demonstrate that the changes in total and paused Pol II binding following aclarubicin treatment are minimal genome-wide.

Signal for Pol II-Ser2p was enriched at 9,968 genes in control samples, indicating their active transcription. In contrast to total Pol II and Pol II-Ser5p, aclarubicin treatment increased Pol II-Ser2p signal at active genes (Fig. 1, C and D). Furthermore, analysis of the difference in Pol II-Ser2p signal between aclarubicin-treated and control samples revealed a net increase at both cCREs and genes (fig. S1, H and I). Collectively, these data indicate that aclarubicin treatment results in an enrichment of the elongating form of Pol II at active genes. While Pol II-Ser2p is traditionally associated with transcriptional elongation, we observed signal localized mainly at the TSS. We attribute this to the Tn5 transposase preferentially targeting accessible chromatin regions, such as TSSs, for tagmentation.

To test whether these observations are unique to aclarubicin or if they apply to other anthracyclines, we performed CUT&Tag targeting total Pol II, Pol II-Ser5p, and Pol II-Ser2p in cells treated for 30 minutes with doxorubicin. Across promoters, doxorubicin minimally affected total Pol II and resulted in a slight decrease in Pol II-Ser5p (fig. S1, J and K). Doxorubicin treatment, similar to aclarubicin treatment, increased Pol II-Ser2p signal at active genes (fig. S1L). We explored the effect of doxorubicin on the distribution of Pol II around the TSS. We found that doxorubicin treatment does not result in substantial changes in Pol II distribution around the TSS (fig. S1, M to O). In contrast, aclarubicin caused gains in total Pol II, Pol II-Ser5p, and Pol II-Ser2p downstream of the TSS. These data suggest that aclarubicin and doxorubicin impact Pol II via distinct mechanisms, particularly within the gene body.

### Aclarubicin redistributes Pol II and substantially accumulates Pol II-Ser2p at histone genes

The histone genes stood out in our genome-wide analyses. These genes are clustered on chromosomes 1 and 6, each organized into phase-separated nuclear compartments called Histone Locus Bodies (HLBs) (*15*). We found these loci were among the most enriched for total and paused Pol II and showed one of the largest aclarubicin-induced gains in elongating Pol II. We therefore sought to understand how aclarubicin impacts Pol II at the histone genes.

We first assessed how aclarubicin treatment changes total Pol II distribution over the histone genes. We generated a plot showing the fold change in total Pol II signal around the TSS of the histone genes (Fig. 2A and fig. S2A). This visualization revealed a decrease in total Pol II signal upstream of the TSS in aclarubicin-treated samples compared with controls. We found this negative fold change in total Pol II signal progressively diminished towards the TSS. Downstream of the TSS, we observed a gain in total Pol II signal.

**Fig. 2.**
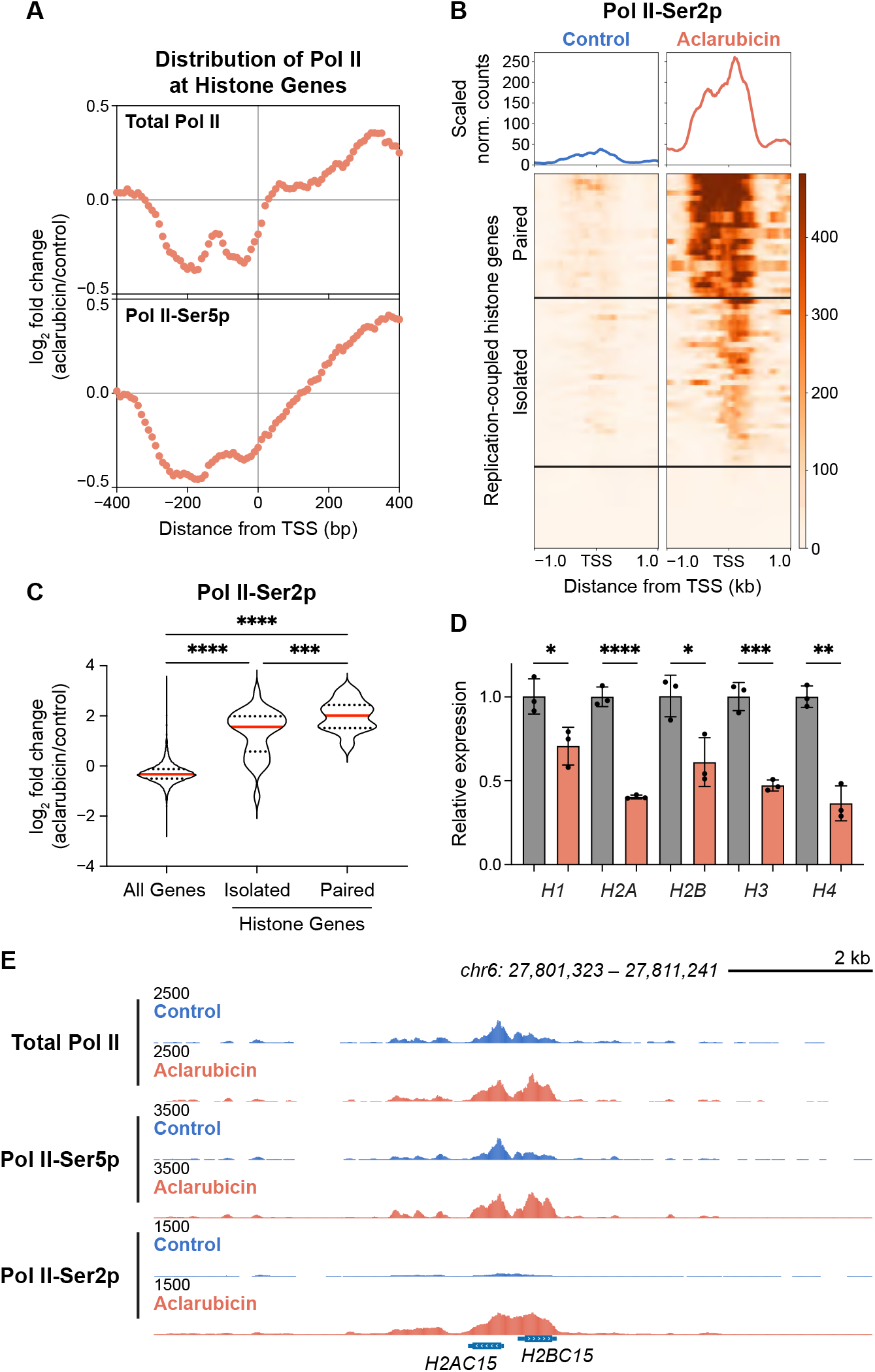
Aclarubicin strongly impacts Pol II distribution and modifications at replication-coupled histone genes. **(A)** Plots showing distribution of fold change (aclarubicin vs. control) in total Pol II and Pol II-Ser5p normalized counts around the TSS of histone genes. **(B)** Average signal distribution plot and heatmap of elongating Pol II (Ser2p) CUT&Tag signal under control and aclarubicin treatment conditions aligned to the TSS of histone genes. **(C)** Violin plots of per-gene log_2_ fold change (aclarubicin vs. control) of Pol II-Ser2p normalized counts across all genes (*n* = 19,305), isolated histone genes (*n* = 43), paired histone genes (*n* = 26). Repeated-measures one-way ANOVA with Geisser-Greenhouse correction and Tukey’s multiple comparisons test was performed on per-replicate median log_2_ fold changes of drug–control pairs (*N* = 8). \*\*\**P* < 0.001, \*\*\*\**P* < 0.0001. Violin plots indicate median, minimum, maximum, and first and third quartiles. **(D)** Histone mRNA levels following aclarubicin treatment (coral) compared with control (grey). Significance was determined by unpaired two-tailed Student’s *t*-test. Bars represent mean ± SD of three biological replicates (*N* = 3). \**P* < 0.05, \*\**P* < 0.01, \*\*\**P* < 0.001, \*\*\*\**P* < 0.0001 versus control. **(E)** Representative genome browser distribution of Pol II under control and aclarubicin treatment conditions at *H2AC15* and *H2BC15* genes.

We found the changes to paused Pol II (Ser5p) following aclarubicin treatment were similar to total Pol II at the histone genes (Fig. 2A and fig. S2B). In visualizing the fold change in Pol II-Ser5p distribution around the TSS following aclarubicin treatment, we observed a decrease in Pol II-Ser5p signal upstream of the TSS. Pol II-Ser5p signal gradually rose towards the TSS and downstream of the TSS. Together, these data demonstrate that aclarubicin redistributes Pol II (total and Ser5p) around the TSS of the histone genes, inducing a loss of polymerase within the promoter region and gain within the gene body (Fig. 2E).

The histone genes are atypical in that they are not enriched for the Ser2p-marked form of Pol II when transcribed (*10, 16*). Accordingly, we observed little Pol II-Ser2p signal across the active histone genes in control samples (Fig. 2B). Strikingly, aclarubicin treatment caused massive gains in Pol II-Ser2p across the histone genes (Fig. 2B and fig. S2C). The histone genes have a dedicated Pol II CTD kinase, cyclin-dependent kinase 11 (CDK11) (*17*). CDK11 phosphorylates serine 2 residues downstream of the TSS within what Gajdušková et al. described as the “transition point” near the middle of the histone genes. CDK11 may be responsible for the gains in Pol II-Ser2p signal. With aclarubicin treatment, Pol II probably transcribes far enough into histone gene bodies to be phosphorylated by CDK11. We found the histone genes that share a divergent promoter region (paired) showed the largest increase in Pol II-Ser2p occupancy (Fig. 2, C and E). Compared with all genes, isolated and paired histone genes showed approximately three- and four-fold increases, respectively, in Pol II-Ser2p signal following aclarubicin treatment. Previous work in *Drosophila* has suggested that divergent promoters trap transcription-induced negative supercoils between them and thus are more sensitive to intercalating drugs (*10*). This phenomenon may account for the effects we observe in human cells.

We quantified Pol II-Ser2p signal at the multiple genes that encode each of the five classes of histone proteins (H1, H2A, H2B, H3, and H4) and found that aclarubicin treatment affects all histone types (fig. S2, D and E). *H1* genes showed the smallest gains in Pol II-Ser2p, while *H2A* and *H2B* genes showed the greatest gains. *H1* is under transcriptional regulation distinct from the octameric histone genes, which may underlie the differences in aclarubicin response. Specifically, the biogenesis factor CRAMP1 binds to *H1* promoter motifs and confers resistance to topoisomerase inhibitors (*18*). Furthermore, in mammals, there is a high frequency of divergently transcribed *H2A–H2B* gene pairs and no divergently transcribed *H3–H4* gene pairs. This feature of *H2A* and *H2B* genes likely contributed to these genes showing the greatest gains in Pol II-Ser2p after aclarubicin treatment. *H2A* and *H2B* genes are also the most highly transcribed histone genes (*16*), indicating that the aclarubicin-induced gain in Pol II-Ser2p is related to transcription rate.

We investigated the potential divergence in the effects of aclarubicin and doxorubicin on Pol II at the histone genes. In doxorubicin-treated samples, we observed subtle decreases in total Pol II and Pol II-Ser5p signal (fig. S2, F and G). We observed an increase in Pol II-Ser2p signal with doxorubicin treatment (fig. S2H). This increase was smaller than the effect observed under aclarubicin conditions. These findings indicate that while doxorubicin and aclarubicin have similar effects on Pol II-Ser2p levels genome-wide, the disproportionate impact on Pol II-Ser2p at histone genes is unique to aclarubicin.

To test if the increase in Pol II-Ser2p at the histone genes results in more histone mRNA production, we measured transcript levels by Reverse Transcription-quantitative PCR (RT-qPCR). Replication-coupled histone gene transcripts lack 3’ polyadenylation processing; thus, we performed RT-qPCR on isolated total RNA with degenerate primers that amplify the multiple genes encoding a given histone class. Histone mRNA production was detected in control samples (Fig. 2D). In aclarubicin-treated samples, after 30 minutes of drug treatment, there was a significant reduction in mRNA abundance for all histone types. These results imply that the gain in Pol II-Ser2p at the histone genes is nonproductive and that aclarubicin treatment results in a rapid reduction in histone mRNA.

### Transcription inhibition synergizes with aclarubicin at histone genes

We were intrigued that aclarubicin resulted in a loss of Pol II upstream of the TSS of the histone genes, as this suggests that aclarubicin intercalation may antagonize Pol II occupancy at promoter regions. To test this possibility, we explored a potential synergistic effect between aclarubicin and triptolide. Triptolide blocks the ATPase activity of general transcription factor TFIIH (*19*). This blockage prevents the melting of promoter DNA such that transcription initiation cannot proceed, triggering the degradation of Pol II (*20*). We treated K562 cells for 1 hour with triptolide, both alone and together with aclarubicin, and performed CUT&Tag as before (Fig. 3A). In the triptolide-only treatment condition, we found a loss of Pol II (total, Ser5p, and Ser2p) signal across promoters (fig. S3, A to C). This effect was more subtle than in other studies (*21*), implying that the treatment here is not as severe. However, this partial triptolide treatment substantially diminished total and paused Pol II levels at the histone genes, indicating that the histone genes are sensitive to triptolide (Fig. 3, B, C, and E). Pol II-Ser2p signal across the histone genes was largely unchanged following triptolide treatment compared with control A and remained at low levels (Fig. 3, D and E). Taken together, these findings highlight the sensitivity of the histone genes to transcriptional perturbation.

**Fig. 3.**
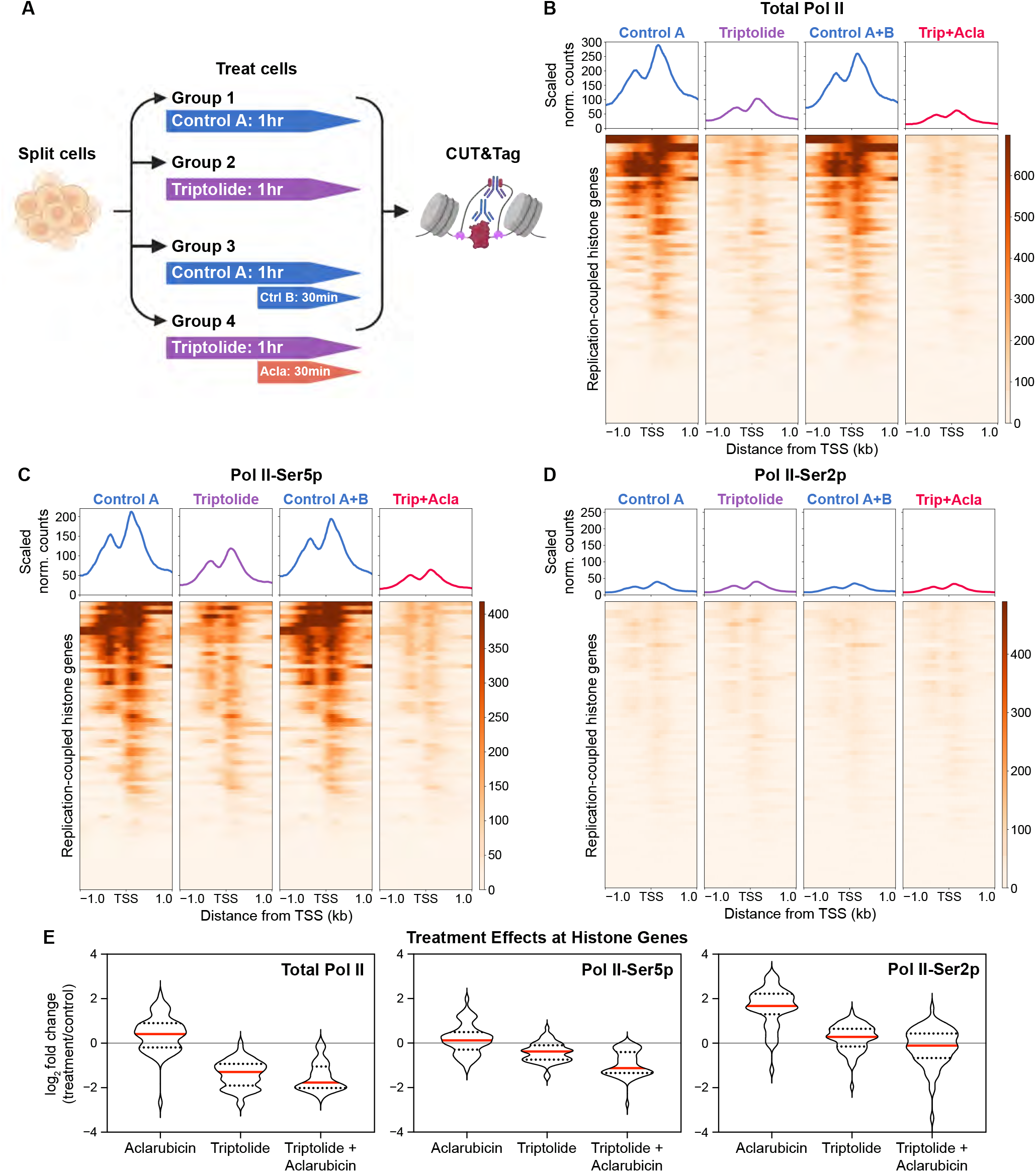
Transcription inhibition synergizes with aclarubicin at histone genes. **(A)** Experimental workflow. K562 cells were split into treatment groups with DMSO (control, ctrl), triptolide (trip), and aclarubicin (acla). Created in BioRender. Nguyen, K. (2026) https://BioRender.com/deebfgs. **(B** to **D)** Average signal distribution plot and heatmap of total Pol II (Rpb1) (B), paused Pol II (Ser5p) (C), and elongating Pol II (Ser2p) (D) CUT&Tag signal under treatment conditions aligned to the TSS of histone genes. For (B) to (D), merged data from three biological replicates per treatment condition shown. **(E)** Violin plots of per-gene log_2_ fold change (treatment vs. control) of Pol II (total, Ser5p, and Ser2p) normalized counts across histone genes. Violin plots indicate median, minimum, maximum, and first and third quartiles.

In the co-treatment condition with triptolide and aclarubicin, we observed similar Pol II occupancy across promoters to that seen in the triptolide-only treatment condition (fig. S3, A to C). At the histone genes, the reduction observed previously for total and paused Pol II in the triptolide-only treatment was further intensified (Fig. 3, B, C, and E), indicating a synergistic effect whereby aclarubicin treatment amplifies the loss of Pol II induced by triptolide. Pol II-Ser2p levels at the histone genes were not affected in the co-treatment condition compared with control A+B (Fig. 3, D and E). This result indicates that in the absence of transcription initiation, aclarubicin treatment does not cause accumulation of elongating Pol II at the histone genes. Collectively, these findings further emphasize the sensitivity of the histone genes to aclarubicin treatment.

### Aclarubicin increases NPAT binding at histone genes

Replication-coupled histone gene transcription is primarily restricted to S phase and requires the Nuclear Protein of the Ataxia Telangiectasia-mutated (NPAT) transcription factor to be recruited to HLBs (*22, 23*). During S phase, the histone genes have the highest density of Pol II across the human genome (*24*). This enrichment is cytologically visible over HLBs (*25*). To visualize histone gene transcription in S-phase cells, we added 5-ethynyl-2′-deoxyuridine (EdU) to K562 cell cultures to label newly replicated DNA, fixed cells, directly conjugated a fluorophore to incorporated EdU, and immunostained for NPAT and Pol II-Ser5p. Of the 519 cells we imaged, 64% (331 cells) were in S phase at the time of EdU treatment, while 36% (188 cells) were in gap phases. We observed bright focal spots of NPAT and Pol II-Ser5p, and their colocalization, in both S-phase and some gap-phase cells (Fig. 4A and fig. S4). The NPAT foci are HLBs, and enrichment of Pol II-Ser5p at HLBs in gap-phase K562 cells has been previously observed (*26*). Corroborating NPAT and Pol II-Ser5p colocalization genomically, we found that the levels of NPAT binding at individual histone genes were tightly correlated with the levels of Pol II-Ser5p at those same genes (Fig. 4B), suggesting that the binding of NPAT and Pol II at the histone genes are coupled.

**Fig. 4.**
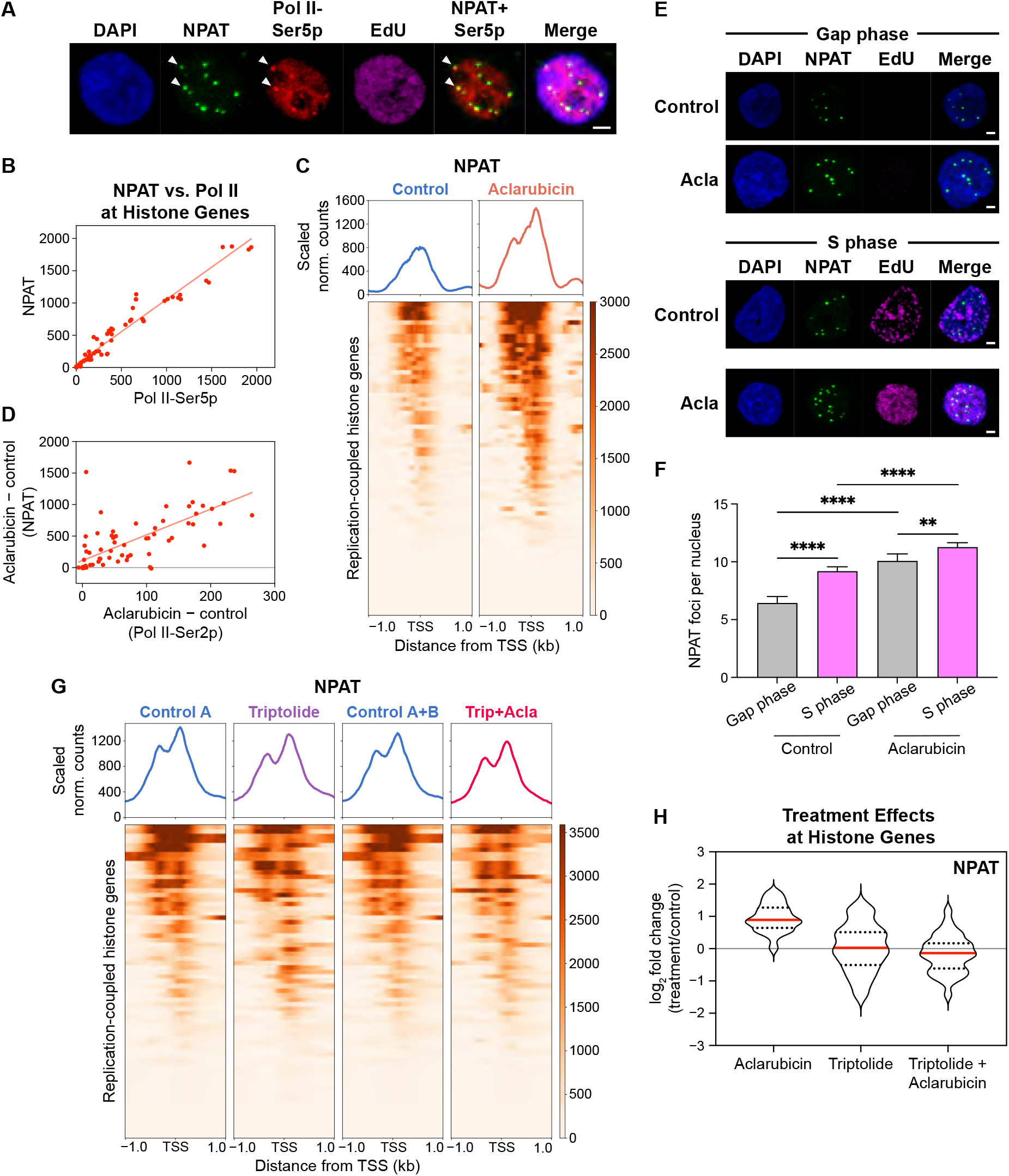
Aclarubicin increases NPAT binding at histone genes. **(A)** Representative immunofluorescence images of S-phase K562 cell treated with vehicle control, stained for DAPI (blue), NPAT (green), Pol II-Ser5p (red), and EdU (magenta). Arrowheads indicate colocalizing NPAT and Pol II-Ser5p foci. Scale bar, 2 µm. **(B)** NPAT versus Pol II-Ser5p normalized counts in control condition at individual histone genes (Spearman ρ = 0.98, \*\*\*\**P* < 0.0001). Simple linear regression line shown for visualization. **(C)** Average signal distribution plot and heatmap of NPAT CUT&Tag signal under control and aclarubicin treatment conditions aligned to the TSS of histone genes. Merged data from three biological replicates per treatment condition shown. **(D)** Scaled aclarubicin-treated (*A*) minus control (*C*) NPAT normalized counts versus (*A* − *C*) Pol II-Ser2p normalized counts at individual histone genes (Spearman ρ = 0.75, \*\*\*\**P* < 0.0001). Simple linear regression line shown for visualization. **(E)** Representative immunofluorescence images of gap-and S-phase cells under each treatment condition. Scale bar, 2 µm. **(F)** Bar graph showing number of NPAT foci per nucleus. Bars represent mean foci count. Error bars indicate 95% confidence intervals. Nuclei count per condition: control gap phase, *n* = 188; control S phase, *n* = 331; aclarubicin-treated gap phase, *n* = 222; aclarubicin-treated S phase, *n* = 373. Welch’s ANOVA and Games-Howell’s multiple comparisons test was performed on foci counts. \*\**P* < 0.01, \*\*\*\**P* < 0.0001. **(G)** Average signal distribution plot and heatmap of NPAT CUT&Tag signal under treatment conditions aligned to the TSS of histone genes. Merged data from three biological replicates per treatment condition shown. **(H)** Violin plots of per-gene log_2_ fold change (treatment vs. control) of scaled NPAT normalized counts across histone genes. Violin plots indicate median, minimum, maximum, and first and third quartiles.

NPAT chromatin binding is highly specific to the histone genes (*27*), and we sought to test whether aclarubicin treatment affects NPAT distribution at these sites. Following aclarubicin treatment, we observed an approximate two-fold increase in NPAT signal over the histone genes (Fig. 4, C and H). At individual histone genes, we found the aclarubicin-induced changes in NPAT levels to be correlated with the changes in Pol II-Ser2p levels at those same sites (Fig. 4D). Collectively, these data suggest that aclarubicin treatment induces an accumulation of transcription machinery at the histone genes.

We investigated whether aclarubicin treatment only affects the histone genes when they are transcriptionally active in S phase of the cell cycle using cytology as before. We found that the aclarubicin-treated and control cell populations contained comparable proportions of EdU-positive cells (63% versus 64%, respectively). Thus, the loss of histone mRNA we observed by RT-qPCR was not due to a cell cycle arrest. We quantified the frequency of NPAT foci and observed that S-phase cells had significantly more NPAT foci per nucleus than gap-phase cells (Fig. 4, E and F). It is likely that the increased binding of NPAT at the histone genes results in more foci detected. Aclarubicin treatment significantly increased NPAT foci detected in both gap- and S-phase cells. The difference in foci between aclarubicin-treated and control samples was larger for cells in gap phase than S phase. Collectively, these data indicate that the enrichment of NPAT binding following aclarubicin treatment occurs independently of the cell cycle.

Given that inhibiting transcription initiation with triptolide alleviated the effect of aclarubicin on Pol II-Ser2p at the histone genes, we examined NPAT binding following the same treatment regimen. NPAT signal across the histone genes was largely unaffected by triptolide treatment compared with control A (Fig. 4, G and H). This indicates that while NPAT levels at individual histone genes were tightly correlated with Pol II-Ser5p, NPAT remains bound to the histone genes once recruited. Interestingly, NPAT signal showed no detectable changes at the histone genes in the co-treatment condition compared with control A+B (Fig. 4, G and H). This lack of response indicates that triptolide treatment abolishes the increased NPAT binding to the histone genes stimulated by aclarubicin treatment. Additionally, this result suggests that the increase in NPAT binding at the histone genes following aclarubicin treatment is driven by Pol II activity.

## Discussion

In this study, we have demonstrated that treatment of human K562 cells with the anthracycline aclarubicin results in accumulation of elongating Pol II (Ser2p) at active genes. This increase was most dramatic at the replication-coupled histone genes. Extending observations originally made in *Drosophila* (*10*), we establish the Histone Locus Body as a site of heightened sensitivity to aclarubicin-mediated chromatin disruption. While the accumulation of Pol II-Ser2p was previously interpreted to indicate more transcription elongation at active genes following aclarubicin treatment (*10*), our measurements showed that productive transcription of the histone genes actually decreases. The gain in the Ser2pmarked form of Pol II at the histone genes after aclarubicin treatment was especially surprising because, in untreated human and *Drosophila* cells, we have measured low levels of Pol II-Ser2p at these genes relative to other genes (*10, 16*). We have also found exceptional enrichment of Pol II-Ser5p at the histone genes relative to the rest of the genome (*10, 16, 28, 29*). These unusual features may result from the reliance of the histone genes on Pol II backtracking at their promoters for regulation instead of promoter-proximal pausing (*26, 30*). The relatively short lengths of the histone genes could also cause this phenomenon, where the time window to deposit the Ser2 phosphorylation mark on the elongating Pol II is narrow. The question remains as to why this modification accumulates at the histone genes after aclarubicin treatment.

Aclarubicin intercalates into DNA at open chromatin regions, including the promoters of active genes (Fig. 5A) (*31–33*). In addition to chromatin accessibility, the torsional landscape of those active promoters also contributes to aclarubicin intercalation. As Pol II elongates into the gene body, it generates negatively supercoiled DNA behind it and positively supercoiled DNA in front (Fig. 5B) (*34*). Aclarubicin intercalates into negative supercoils with high affinity (*35*). Thus, transcription progression allows aclarubicin to intercalate beyond the promoter region, and the high transcriptional flux of Pol II at the histone genes likely facilitates greater aclarubicin intercalation (Fig. 5, C and D). Aclarubicin inhibits topoisomerases (*36–39*). As positive supercoils accumulate ahead of the advancing polymerase, torsional stress accumulates and is unrelieved. Without topoisomerases present to remove this torsion, eventually, Pol II translocation is inhibited (*40, 41*). Therefore, it is plausible that genes enriched with aclarubicin become more difficult to transcribe and, over time, Pol II progression slows down or stalls. In this way, the accumulation of Pol II-Ser2p at the histone genes may be due to increased Pol II dwell time within histone gene bodies, allowing time for kinases to phosphorylate the CTD at Ser2 residues (*42*). These aclarubicin effects appear to occur genome-wide as Pol II-Ser2p is elevated across all active genes.

**Fig. 5.**
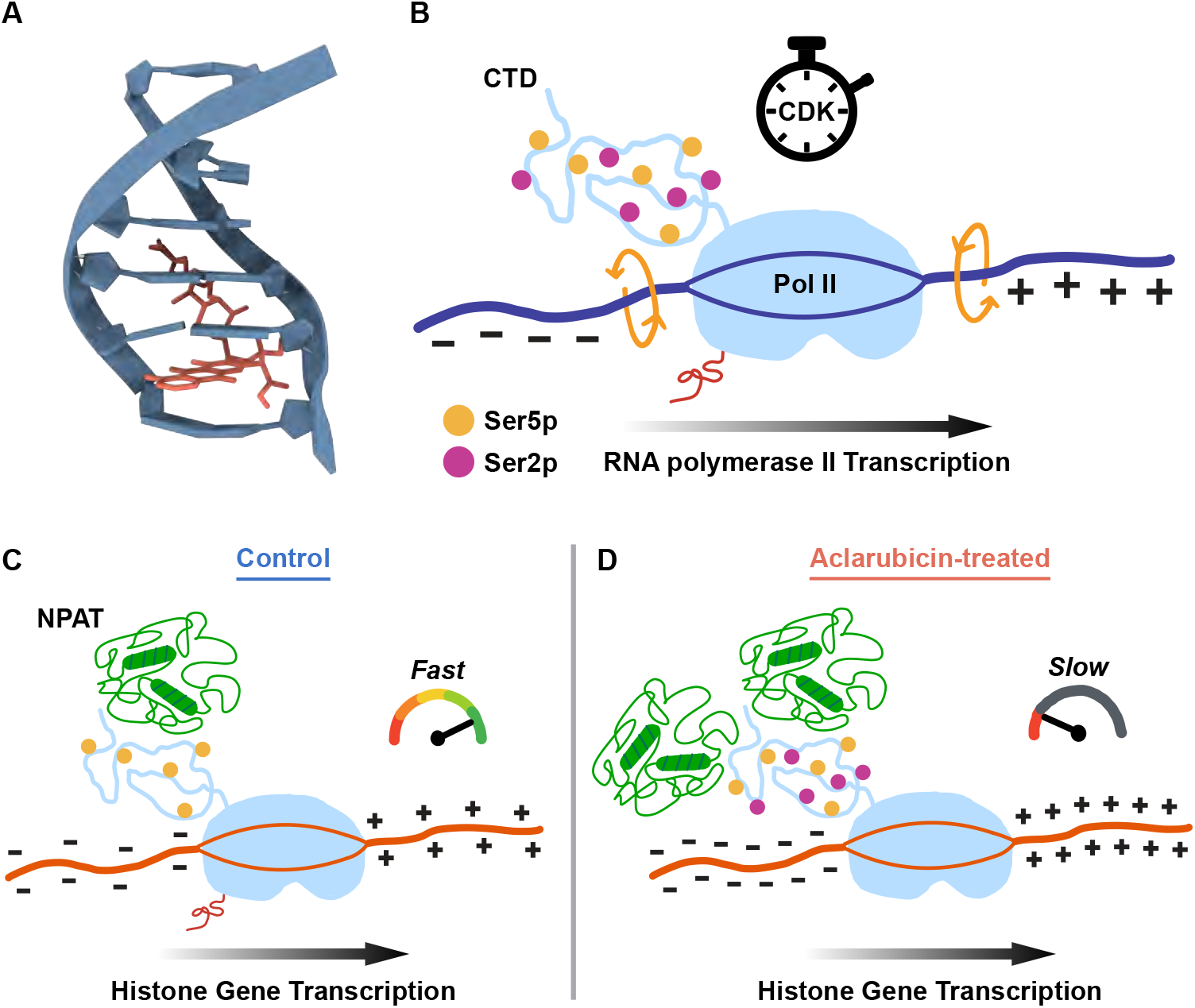
Impact of aclarubicin intercalation on Pol II transcription. **(A)** Aclarubicin–DNA complex structure. Aclarubicin intercalates into DNA and separates adjacent base pairs. This process unwinds the DNA helix locally and, in a topologically constrained context, results in positive supercoiling. The trisaccharide sugar moiety of aclarubicin sits inside the DNA minor groove. **(B)** Topological representation of Pol II transcription. Pol II translocation is controlled by a balance of torque, where Pol II rotation is counterbalanced by DNA twisting that generates negatively supercoiled domains upstream and positively supercoiled domains downstream of the enzyme (twin-supercoiled-domain model). Pol II speed is determined by torsion in front and is kinetically coupled with kinase activity, as represented by the stopwatch. Ser5p and Ser2p marks are not drawn to scale or frequency on the Pol II CTD. Nascent messenger RNA is indicated with red line exiting transcribing Pol II. Negatively (−) and positively (+) supercoiled DNA are indicated. **(C)** Histone gene transcription requires NPAT. Pol II transcription at the histone genes is rapid and abundant, generating high levels of negative and positive DNA supercoiling that demands increased activity of topoisomerases to relieve. Histone gene transcription occurs without Pol II-Ser2p. **(D)** Aclarubicin intercalation at the histone genes is ubiquitous, resulting in the loss of topoisomerase activity and accumulation of positive supercoils ahead of advancing polymerases. This torsional barrier slows or stalls Pol II progression, allowing time for Ser2 phosphorylation of the Pol II CTD at the histone genes. NPAT responds to the altered transcriptional status of Pol II at the histone genes and increases in abundance.

Aclarubicin treatment has been shown to broadly impact chromatin structure (*4, 10*). Therefore, due to torsional effects, aclarubicin may be promoting accessible chromatin regions beyond the histone genes. While sites of chromatin accessibility are likely coupled with transcription initiation (*43*), we did not observe a change in Pol II-Ser5p signal in the promoter region of active genes. However, at the histone genes, we observed a decrease in Pol II-Ser5p within the promoter region. This could be explained by two possible mechanisms. First, the negatively supercoiled DNA produced behind advancing polymerases at promoters is filled with intercalated aclarubicin. The presence of aclarubicin would preclude Pol II binding at these sites. This process is intensified at the histone genes. The histone loci are regions full of actively engaged Pol II and under intense transcriptional demand (Fig. 5C). As such, histone gene promoters, especially those that are divergent, are prone to accumulating negative supercoils and, consequently, aclarubicin. Cyclically, the inhibition of topoisomerase activity by aclarubicin further exacerbates the levels of negative DNA supercoiling and promotes its own intercalation.

Second, promoter melting during transcription initiation exposes intercalation sites for aclarubicin. In a topologically constrained context, promoter melting and DNA helix unwinding accompanying aclarubicin intercalation (*44*) generate positive DNA supercoils that propagate outward from the promoter. These positive supercoils unwrap flanking nucleosomes. Enhanced nucleosome unwrapping increases nucleosome turnover (*8*). Alterations to the chromatin landscape in this regard are permissive of transcription, especially destabilization of the +1 nucleosome, which is the first nucleosome downstream of the TSS that is a barrier to transcription elongation. In the context of aclarubicin treatment, this destabilization could further promote the Pol II already loaded to move into the gene body. This is evident at regions with high transcriptional flux, such as the histone genes. After inhibiting transcription initiation with triptolide, we observed a decrease in Pol II-Ser5p at the histone genes. Co-treating with aclarubicin augmented this drop further. These findings suggest that the reduced elongation barrier in the context of aclarubicin treatment allows loaded Pol II to rapidly release downstream at a greater rate, depleting the Pol II-Ser5p pool ***Slow*** faster at these high-flux genes.

What underlying mechanism drives the increase in NPAT abundance at the histone genes following aclarubicin treatment? These gains could be associated with changes in Pol II, in particular its conversion from initiating to elongating (Fig. 5D). Our study indicates that NPAT responds to the transcriptional status of the histone genes to adjust NPAT levels. This adjustment must involve Pol II, as triptolide treatment blocked the NPAT increase induced by aclarubicin. A recent study in *Drosophila* by Kemp et al. demonstrated that depletion of Pol II attenuates HLB maturation and reduces levels of Mxc, the *Drosophila* counterpart of human NPAT (*45*). Kemp et al. also showed that the transition of Pol II from paused to elongating is a major cell-cycle-regulated step that couples histone transcription to S phase. These findings support the interdependence between Pol II and HLB activity.

Extending a model from a recent live-cell imaging study offers complementary support for our proposed aclarubicin-induced increase in Pol II dwell time. Abidi et al. showed that intrinsically disordered regions on transcription factors can mediate heterotypic interactions and locus-specific enrichment through transient contacts independent of a structured DNA-binding domain (*46*). NPAT does not have a DNA-binding domain and is predicted to be nearly 90% disordered according to the Predictor of Natural Disordered Regions (PONDR) (*47*). It is well-documented that the Pol II CTD, which also has an intrinsically disordered structure (*48*), recruits transcription-associated proteins (*49, 50*). Since NPAT is almost entirely disordered, it presents a large surface of weak contact sites. These contact sites would support extensive multivalent interactions if NPAT engages the Pol II CTD (Fig. 5, C and D). Therefore, the increased NPAT binding at the histone genes that we observed following aclarubicin treatment would raise the local concentration of NPAT–CTD contact sites. While each NPAT–CTD interaction would be transient, the high availability of NPAT could favor rapid Pol II reengagement after dissociation, thus lengthening the apparent residence time of Pol II at the histone genes through an avidity-based mechanism.

Our observations at the histone genes appear to be specific to aclarubicin and not to other anthracyclines, such as doxorubicin. The differences in chemical structure between aclarubicin and doxorubicin may underlie their differing impacts on Pol II activity. Doxorubicin intercalation leads to the formation of covalent DNA adducts (*51*), which directly antagonize Pol II elongation (*52*). Aclarubicin, however, lacks the chemical residues predicted to form covalent attachments, therefore allowing Pol II to elongate into the gene body. Additionally, doxorubicin and aclarubicin have differing impacts on topoisomerase. Doxorubicin poisons topoisomerase II by covalently trapping the enzyme to cleaved DNA (*53*), resulting in DNA breaks. Aclarubicin inhibits topoisomerase II by preventing the conformational changes required for the enzyme to bind DNA. Aclarubicin also inhibits topoisomerase I. Therefore, the effects of aclarubicin on the histone genes are reversible, but essentially irreversible for doxorubicin.

We have recently shown that histone gene overexpression predicts aggressiveness in human cancer (*28*). Thus, it is of particular oncological interest that aclarubicin effects are most severe at the histone genes. While the effectiveness of anthracyclines in solid tumors is more variable than in liquid tumors, possibly due to limited tissue penetration, an active area of research involves modifying the chemical properties of anthracyclines such that they are more therapeutically accessible (*54*). Structure–activity analyses of anthracyclines have indicated that chemical modifications enhancing chromatin damage correlate with drug potency (*7, 55, 56*). This may give an advantage to aclarubicin treatments over the current standard of care— doxorubicin—which has a weaker effect on chromatin and a DNA-damaging effect (*51*). A purely chromatin-based anticancer mechanism would mitigate the off-target effects of DNA damage, such as secondary cancer mutations and cardiotoxicity (*4*). Two anticancer inhibitors are in clinical trials that have acute transcriptional consequences that converge on the replication-coupled histone genes and HLB. These are inhibitors of protein arginine methyltransferase PRMT5 (*57*) and protein kinase ATR (*58*), master regulator of the DNA damage response pathway. This convergence likely reflects the sensitivity of the histone genes, which operate under unique regulation, and supports the possibility that the antiproliferative efficacy of these small molecules stems largely from histone gene dysregulation.

## Materials and Methods

### Human cell culture and drug treatment

Human K562 cells (ATCC, CCL-243) were cultured in Iscove’s Modified Dulbecco’s Medium (IMDM, ATCC, 30-2005) supplemented with 10% heat-inactivated fetal bovine serum (HI-FBS, HyClone, SH30070.03). Cells were cultured in suspension at 37°C in a humidified incubator set to 5% CO_2_ and were passaged every 2 days. Cell characteristics were monitored and obtained using a Vi-CELL BLU Cell Viability Analyzer (Beckman Coulter). Aclarubicin (Cayman Chemical Company, 15993), doxorubicin (Sigma-Aldrich, D1515), and triptolide (Selleckchem, S3604) used in this study were resuspended in dimethyl sulfoxide (DMSO) and frozen in aliquots. Aclarubicin, doxorubicin, and triptolide aliquots were at 10 mM concentration each. Aclarubicin and doxorubicin were added to cell culture media at a final concentration of 1 µM. Triptolide was added to cell culture media at a final concentration of 10 µM. Cells were also treated with DMSO (1:10000, v/v, for aclarubicin and doxorubicin; 1:1000, v/v, for triptolide) and harvested in parallel with each treatment condition as a control. For CUT&Tag profiling, cells treated with aclarubicin and doxorubicin were harvested after 30 minutes, while cells treated with triptolide were harvested after 1 hour. For imaging and RTqPCR, cells were treated with aclarubicin for 30 minutes prior to harvest. To assess the effects of aclarubicin on cellular characteristics, cells were harvested at 30 minutes, 4 hours, or 24 hours post-treatment.

### Antibodies

For CUT&Tag, we used the following primary antibodies: rabbit monoclonal anti-Phospho-Ser2-Rpb1 CTD (Cell Signaling Technologies, 13499), rabbit monoclonal anti-Phospho-Ser5-Rpb1 CTD (Cell Signaling Technologies, 13523), mouse monoclonal anti-RNA Polymerase II 8WG16 (Rpb1) (Covance, MMS-126R), mouse monoclonal anti-NPAT (Abcam, ab307837). To assist in Tn5 tethering, we also employed the following secondary antibodies: guinea pig polyclonal anti-rabbit IgG (Antibodies Online, ABIN101961), rabbit polyclonal anti-mouse IgG (Abcam, ab46540). We used the following antibodies for immunofluorescence imaging: rabbit monoclonal anti-Phospho-Ser5-Rpb1 CTD (Cell Signaling Technologies, 13523), mouse monoclonal anti-NPAT (Abcam, ab307837), goat polyclonal anti-rabbit Rhodamine Red-X (RRX) (Jackson ImmunoResearch, 111-297-003), goat polyclonal anti-mouse Alexa Fluor 488 (Jackson ImmunoResearch, 115-547-003). Final concentrations for each antibody are specified below.

### CUT&Tag chromatin profiling

On day 1, CUT&Tag was performed as previously described (*13, 21*) with slight modifications. In summary, whole K562 cells were harvested from culture, spun down at 350 X g for 3 minutes, resuspended in wash buffer (23.75 mL ddH_2_O, 500 µL 1M HEPES-NaOH pH 7.5 [20 mM], 750 µL 5M NaCl [150 mM], 6.25 µL 2M spermidine [0.5 mM], 1 Roche cOmplete mini, EDTA-free Protease Inhibitor tablet), and placed on ice. BioMag Plus Concanavalin A (Con A)-coated magnetic beads (Bangs Laboratories, BP531) were activated in binding buffer (9.68 mL ddH_2_O, 200 µL 1M HEPES-KOH pH 7.9 [20 mM], 100 µL 1M KCl [10 mM], 10 µL 1M CaCl_2_ [1 mM], 10 µL 1M MnCl_2_ [1 mM]) as follows. 2 µL of Con A bead slurry per reaction was combined with 20 µL of bead binding buffer per reaction and mixed. Beads were washed twice before resuspending in 20 µL of binding buffer per reaction as before. Cells were counted using Vi-CELL and 50,000 cells per reaction were combined with 20 µL of activated bead suspension and let bind for 10 minutes at room temperature. After binding, beads were never mixed, resuspended, or vortexed to minimize loss. Triton-wash buffer (47 mL ddH_2_O, 1 mL 1M HEPES-NaOH pH 7.5 [20 mM], 1.5 mL 5M NaCl [150 mM], 12.5 µL 2M spermidine [0.5 mM], 500 µL of 10% Triton X-100 [0.1%], 1 Roche cOmplete, EDTAfree Protease Inhibitor tablet) was prepared. Then, cells were lightly cross-linked in 0.1% formaldehyde on beads for 2 minutes as before (*59*), quenched, washed, and brought up in a 1:50 dilution of primary antibody in primary antibody buffer (1 mL triton-wash buffer, 4 µL 0.5M EDTA [2 mM], 4 µL of 30% Bovine Serum Albumin solution [0.1%] (BSA, Sigma-Aldrich, A8577)). Primary antibody incubation was performed overnight at 4°C.

On day 2, primary antibody solution was removed, cells were brought up in a 1:100 dilution of secondary antibody solution in triton-wash buffer and were incubated on a rotator at room temperature for 1 hour. Triton-300-wash buffer (24.3 mL triton-wash buffer, 750 µL 5M NaCl [300 mM]) was prepared. Cells were washed with triton-wash buffer, brought up in a 1:20 dilution of pAG-Tn5 (EpiCypher, 15-1117) in triton-300-wash buffer, and incubated on a rotator at room temperature for 1 hour. After, cells were washed with triton-300-wash buffer, resuspended in tagmentation buffer (780 µL ddH_2_O, 200 µL N,N-dimethylformamide (DMF) [20%], 10 µL 1M TAPS pH 8.5 [10 mM], 5 µL 1M MgCl_2_ [5 mM]), and incubated in a PCR cycler at 55°C with a heated lid for 1 hour to complete tagmentation reaction. Cells were then washed in TAPS wash buffer (1 mL ddH_2_O, 10 µL 1M TAPS pH 8.5 [10 mM], 0.4 µL 0.5M EDTA [0.2 mM]) and resuspended in 5 µL of 1% SDS-ProtK release solution (178 µL ddH_2_O, 20 µL of 10% SDS [1%], 2 µL 1M TAPS pH 8.5 [10 mM], 20 µL Thermolabile Proteinase K (NEB, P8111S)). Cells were then incubated in a PCR cycler with a heated lid at 37°C for 1 hour to release Tn5 from tagmented DNA and then 58°C for 1 hour to inactivate Proteinase K. Then, SDS was quenched by adding 15 µL of 6% Triton-X100, vortexing vigorously, and separating supernatant from beads to fresh tubes. DNA was amplified by PCR with 2 µL each of uniquely barcoded 10 µM i5 and i7 primers pairs and 25 µL of NEBNext High-Fidelity 2X PCR Master Mix (NEB, M0541L). The following PCR cycling program was used: 58°C for 5 minutes, 72°C for 5 minutes, 98°C for 5 minutes, 14 cycles of (98°C for 10 seconds, 60°C for 10 seconds), 72°C for 1 minute, hold at 8°C. After, these CUT&Tag libraries were cleaned with 1.3X volume of HighPrep PCR Purification System (MagBio, AC-60500), washed twice with 80% ethanol, resuspended in 27 µL 10 mM Tris-HCl pH 8, let sit for 10 minutes, and extracted to fresh tubes. Libraries were analyzed on an Agilent 4200 TapeStation and pooled at equal volumes for sequencing.

### DNA sequencing and data processing

CUT&Tag libraries were sequenced on an Illumina NovaSeq X using paired-end 50 bp reads. Adapters were trimmed and reads shorter than 20 bp were discarded by Cutadapt v4.4 (*60*):

~~~
-j 8 --nextseq-trim 20 -m 20 -a AGATCGGAA-
GAGCACACGTCTGAACTCCAGTCA -A AGATCGGAA-
GAGCGTCGTGTAGGGAAAGAGTGT –Z
~~~

Reads were aligned to the hg19 *Homo sapiens* reference genome obtained from UCSC using Bowtie2 v2.5.1 (*61*):

~~~
--very-sensitive-local --soft-clipped-un-
mapped-tlen --dovetail --no-mixed
--no-discordant -q --phred33 -I 10 -X 1000
~~~

### Data analysis and visualization

Properly paired reads were extracted from the hg19 alignments using bedtools v2.30.0 (`bamtobed -bedpe`) (*62*) and put into BED files of mapped fragments. Bedtools v2.30.0 (`genomecov`) was used to generate coverage-normalized bigWig files. At each base pair, the fraction of counts was normalized by the size of the hg19 reference genome (3,095,693,983 bp) such that a uniform distribution of reads would yield a value of one at each position. Next, scaling factors to calibrate read counts between control and drug-treated samples were generated by summing the total mapped fragments across replicates for each condition. As before (*63*), the scaling factor was calculated as the summed mapped fragments of the drug-treated condition divided by the summed mapped fragments of the paired control condition. The resulting scaling factor was applied to the merged drug-treated bigWig using deepTools v3.5.4 (`bigwigCompare`) (*64*) with the merged control bigWig as reference. Scaling factors were uniquely generated for each epitope and only applied within replicate-matched drug–control comparisons.

We prepared several gene annotation files for our downstream analyses. For Candidate *cis*-Regulatory Elements (cCREs), we downloaded the hg38 version from ENCODE (*14*) and re-positioned coordinates to hg19 using UCSC’s LiftOver tool (*65*), resulting in 2,343,081 entries. Given that many cCREs were located in repetitive regions of the genome, we intersected the hg19 cCRE file with UCSC’s RepeatMasker regions using Bedtools v2.30.0 (`intersect -v`) to remove such entries, resulting in a BED file of 1,036,067 entries. For gene-level analyses with Bland-Altman plots, heatmaps, and fold-change quantification we used a non-redundant gene list of 19,305 protein-coding genes obtained from NCBI for hg19 (table S1), from which lists of histone genes were curated (table S2). Human replication-coupled histone genes were annotated as described by Seal et al. (*66*) and filtered to just those clustered on chromosomes 1 and 6 (*28*). For total Pol II (Rpb1), Pol II-Ser5p, and NPAT, we defined our gene window as 500 bps upstream and downstream of the TSS. For Pol II-Ser2p, we defined our gene window as 200 bps upstream of the TSS in addition to the entire length of the gene body.

Bland-Altman plots (*67*) were generated to compare drug-treated versus control signal across cCREs and genes. To do this, we used Bedtools v2.30.0 (`intersect`) and (`groupby`) to intersect merged, genome coverage-normalized bedGraph files with cCREs or gene BED files and sum counts within each region. We further controlled for differences in region size by dividing the summed values by the length of each cCRE region or base pair window, resulting in mean per-base-pair signal per cCRE or gene window. Paired values from aclarubicin-treated (*A*) and control (*C*) samples were plotted as aclarubicin minus control (*A* − *C*) versus log_10_ (*A* + *C*)/2.

Heatmaps were generated using the merged, scalingfactor-calibrated bigWig files for drug-treated and control conditions as input. Signal counts matrices were first computed with deepTools v3.5.4 (`computeMatrix`) and then visualized with (`plotHeatmap`).

For quantification of signal changes at regions following drug treatment, we used deepTools v3.5.4 (`multiBigwigSummary`) with individual replicate, unscaled genome coverage-normalized bigWig files as input. A rank-ordered knee plot of replicate-averaged control normalized counts across all genes was used to identify signal threshold. Regions were retained if the control signal exceeded this threshold. For violin plots, per-gene log_2_ fold changes were calculated as the log_2_ ratio of mean drug versus mean control signal across replicates. For statistical tests, per-gene log_2_ fold change was computed for each matched replicate pair as log_2_ ((drug + *ε*) / (control + *ε*)), where *ε* = half the smallest nonzero value across all genes and replicates. Afterward, median log_2_ fold changes across genes were computed within each replicate pair.

Representative genome tracks from merged, scalingfactor-calibrated bigWig files were visualized using UCSC’s Genome Browser.

### RNA extraction and cDNA synthesis

Following treatment, samples of 200,000 K562 cells were harvested, with three biological replicates per condition (control and aclarubicin-treated). Cells were spun down at 350 X g for 3 minutes and total RNA extraction was performed using the Macherey-Nagel NucleoSpin RNA mini kit for RNA purification (Takara Bio, 740955) following manufacturer’s instructions with 10 mM of reducing agent dithiothreitol (DTT, Thermo Scientific, P2325) used as an alternative to beta-mercaptoethanol (BME) within the Lysis Buffer RA1. Eluted RNA was combined with 1 unit of RNasin Ribonuclease Inhibitor (Promega, N2515) per µL and quantified by spectrophotometry (Thermo Scientific, ND-ONE-W). RNA integrity was assessed via an Agilent 4200 TapeStation. cDNA was synthesized using Moloney Murine Leukemia Virus Reverse Transcriptase (M-MLV RT, Invitrogen, 28025013) following manufacturer’s instructions with a 20 µL reaction volume containing 1 µg of total RNA, 1 µL 50 µM random hexamers [2.5 µM] (Invitrogen, N8080127), 1 µL 10 mM dNTP mix [0.5 mM] (Kapa Biosystems, KN1009), 4 µL 5X First-Strand Buffer [1X] (Invitrogen, 28025013), 2 µL 100 mM DTT [10 mM] (Invitrogen, 28025013), 1 µL 40 U/µL RNasin Ribonuclease Inhibitor [2 U/µL] (Promega, N2515), and nuclease-free H_2_O up to 20 µL. After the RT reaction and inactivation, RNA complementary to cDNA was removed by adding 0.5 µL 5,000 U/mL RNase H [0.125 U/µL] (NEB, M0297) followed by an incubation in a PCR cycler at 37°C with a heated lid for 20 minutes.

### Histone gene qPCR and data analysis

Production of cDNA was assumed to be equivalent to input RNA (1 µg). Each reaction contained 1 ng cDNA, 10 µL 2X SYBR Green PCR Master Mix [1X], (Applied Biosystems, 4309155), 3.2 µL target primer mix with both 10 µM forward and reverse primers [1.6 µM], and nucleasefree H_2_O up to 20 µL. Primer sequences for histone genes were as previously described by Paranal et al. (*68*). Quantitative PCR was performed on a 384-well QuantStudio 5 Real-Time PCR System (Applied Biosystems) using the following cycling program: 50°C for 2 minutes, 95°C for 5 minutes, 40 cycles of (95°C for 15 seconds, 58°C for 30 seconds), melt curve: 95°C for 15 seconds, ramp down at 1.6°C/s to 60°C for 1 minute, ramp up at 0.075°C/s to 95°C for 15 seconds with continuous fluorescence acquisition. Three biological replicates were analyzed per condition with each replicate measured for all primer sets in technical quadruplicate. After cycling, Ct values, amplification plots, and derivative melt curves were automatically generated by the QuantStudio Design and Analysis Software v1.4.3. The mean Ct value of technical quadruplicates was used for analysis.

Fold change in gene expression was calculated as follows. For each biological replicate, a single reference Ct value was generated by taking the geometric mean of the two reference genes (*GAPDH* and *ACTB*). Afterwards, the ΔCt for each target was calculated by subtracting the reference Ct from the target Ct within the same replicate, resulting in three ΔCt values per condition. We then defined a calibrator as the mean ΔCt of the three control replicates. Then, ΔΔCt was calculated by subtracting the calibrator from each ΔCt value for every replicate across both conditions. Subsequently, relative expression was calculated as 2^-ΔΔCt^. Statistical differences were assessed by unpaired two-tailed Student’s *t*-test for each target.

### Immunofluorescence labeling

On day 1, 10 mM 5-ethynyl-2′-deoxyuridine (EdU, Thermo Fisher Scientific, C10640) was added to cell culture media at a final concentration of 10 µM and incubated for 20 minutes. Samples of 100,000 cells were harvested, spundown at 350 X g for 3 minutes, and resuspended in 500 µL ice-cold 1X phosphate-buffered saline (PBS) with 4% formaldehyde. Fixation occurred on ice for 5 minutes, after which the reaction was quenched by adding 25 µL of 2.5M glycine (125 mM). Cells were spun down and resuspended in 1 mL ice-cold PBS with 0.1% Triton X-100 (PBST). Afterward, cells were spun down onto glass slides (Fisherbrand, 12-550-15) using a Cytospin 4 Centrifuge (Thermo Scientific). Slides were washed with PBS in a Coplin jar, placed in a humid chamber, and incubated with wash buffer (42 mL ddH_2_O, 1 mL 1M HEPES-NaOH pH 7.5 [20 mM], 1.5 mL 5M NaCl [150 mM], 12.5 µL 2M spermidine [0.5 mM], 500 µL of 10% Triton X-100 [0.1%], 0.25 g Bovine Serum Albumin [0.5%] (BSA, Sigma-Aldrich, A9647), 5 mL of 5% Casein [0.5%] (Sigma-Aldrich, C7078), 1 Roche cOmplete, EDTA-free Protease Inhibitor tablet) for 30 minutes of blocking at room temperature. In parallel, primary antibodies were diluted 1:200 in antibody buffer (1 mL wash buffer, 4 µL 0.5M EDTA [2 mM]) and pre-blocked for 30 minutes at 4°C. After, 20 µL of the preblocked primary antibodies were added to each slide and slides were placed in a humid chamber at 4°C overnight. On day 2, slides were washed with PBS in a Coplin jar and blocked again with wash buffer for 30 minutes at room temperature. In parallel, secondary antibodies were diluted 1:500 in wash buffer and pre-blocked for 30 minutes at 4°C. After, 20 µL of the pre-blocked secondary antibodies were added to each slide and slides were placed in a humid chamber at 4°C for 1 hour. Next, slides were washed with PBS and EdU was detected using the Click-iT Plus (Thermo Fisher Scientific, C10640) reaction cocktail per manufacturer’s instructions (1X Click-iT reaction buffer, copper protectant, Alexa Fluor 647 picolyl azide, and 1X reaction buffer additive, with ingredients mixed in this order). Slides were incubated with 100 µL of reaction cocktail for 1 hour at room temperature, protected from light. After, slides were washed with fresh PBS twice and stained with 0.5 µg/mL DAPI in PBS for 5 minutes at room temperature, protected from light. Finally, slides were washed with PBS, mounted with ProLong Glass Antifade Mountant (Thermo Fisher Scientific, P36984), and cured overnight prior to imaging.

### Immunofluorescence imaging and analysis

All imaging was performed on a Leica Stellaris 8 laser scanning confocal microscope using a Leica HC PL APO CS2 63x/1.40 oil immersion objective. DAPI-stained DNA, Alexa Fluor 488 (NPAT), Rhodamine Red-X (Pol II-Ser5p), and Alexa Fluor 647 (EdU) were excited at 405, 488, 561, and 638 nm, respectively. Emission was collected sequentially on Leica HyD detectors: 422–501 nm (DAPI), 501–567 nm (Alexa Fluor 488, NPAT), 583–651 nm (Rhodamine Red-X, Pol II-Ser5p), and 651–792 nm (Alexa Fluor 647, EdU). LAS X v4.9.1.30715 software was used to acquire images as 14-slice z-stacks (0.3 µm step size, 1024 × 1024 pixels, 0.18 µm pixel size, 12-bit depth). Identical acquisition settings were used across all conditions. For each four-channel image series, pseudocolored maximum intensity z-projections were generated per channel and split into individual channel images. Images were saved for downstream analysis and assembly in Adobe Illustrator. Nuclei were segmented from the DAPI channel and S-phase nuclei were identified by EdU incorporation.

### Molecular modeling of aclarubicin–DNA interaction

Aclarubicin intercalation within DNA was modeled using a template-based docking approach. Assembly and visualization occurred within UCSF ChimeraX (*69*). The crystal structure of a doxorubicin–DNA complex (*70*) was obtained from the RCSB Protein Data Bank (*71*) and used as the receptor template. Intercalated doxorubicin was removed and replaced by a 3D model of the aclarubicin lig- and created from its isomeric SMILES obtained from Pub-Chem (*72*). Molecular docking was performed with Auto-Dock Vina v1.2.7 (*73, 74*).

## Acknowledgements

We are grateful to our colleagues at the Fred Hutch Cancer Center, previous and current members of the Henikoff lab, and S. Yoo for helpful discussions and feedback on the manuscript. We thank J. Henikoff, C. Codomo, and D. Xu for technical assistance.

## Author contributions

Conceptualization: K.K.N., M.W., and S.H.

Methodology: K.K.N. and M.W.

Investigation: K.K.N. and M.W.

Data curation: K.K.N. and M.W.

Formal analysis: K.K.N. and M.W.

Validation: K.K.N. and M.W.

Visualization: K.K.N.

Resources: M.W. and S.H.

Funding acquisition: M.W. and S.H.

Supervision: M.W. and S.H.

Project administration: M.W. and S.H.

Writing—original draft: K.K.N.

Writing—review & editing: K.K.N., M.W., K.A., and S.H.

## Data, code, and materials availability

All data and code needed to evaluate and reproduce the conclusions in the paper are present in the paper and/or the Supplementary Materials. All raw and processed sequencing data generated in this study have been deposited in the NCBI Gene Expression Omnibus (GEO; https://www.ncbi.nlm.nih.gov/geo/) under accession number GSE334543. Custom scripts used in this study are available from Zenodo (https://doi.org/10.5281/zenodo.20571836). This study did not generate new materials.

## Supplementary materials

Figs. S1 to S4 Tables S1 to S3

## Supplementary Information

**Table S1: Table S1.xlsx**, excel file containing annotated human gene coordinates from Hg19 genome assembly.*

**Table S2: Table S2.xlsx**, excel file containing human replication-coupled histone gene coordinates.*

**Table S3: Tabel S3.xlsx**, excel file containing control oligonucleotides used in this study for RT-qPCR.*

**Fig. S1.**
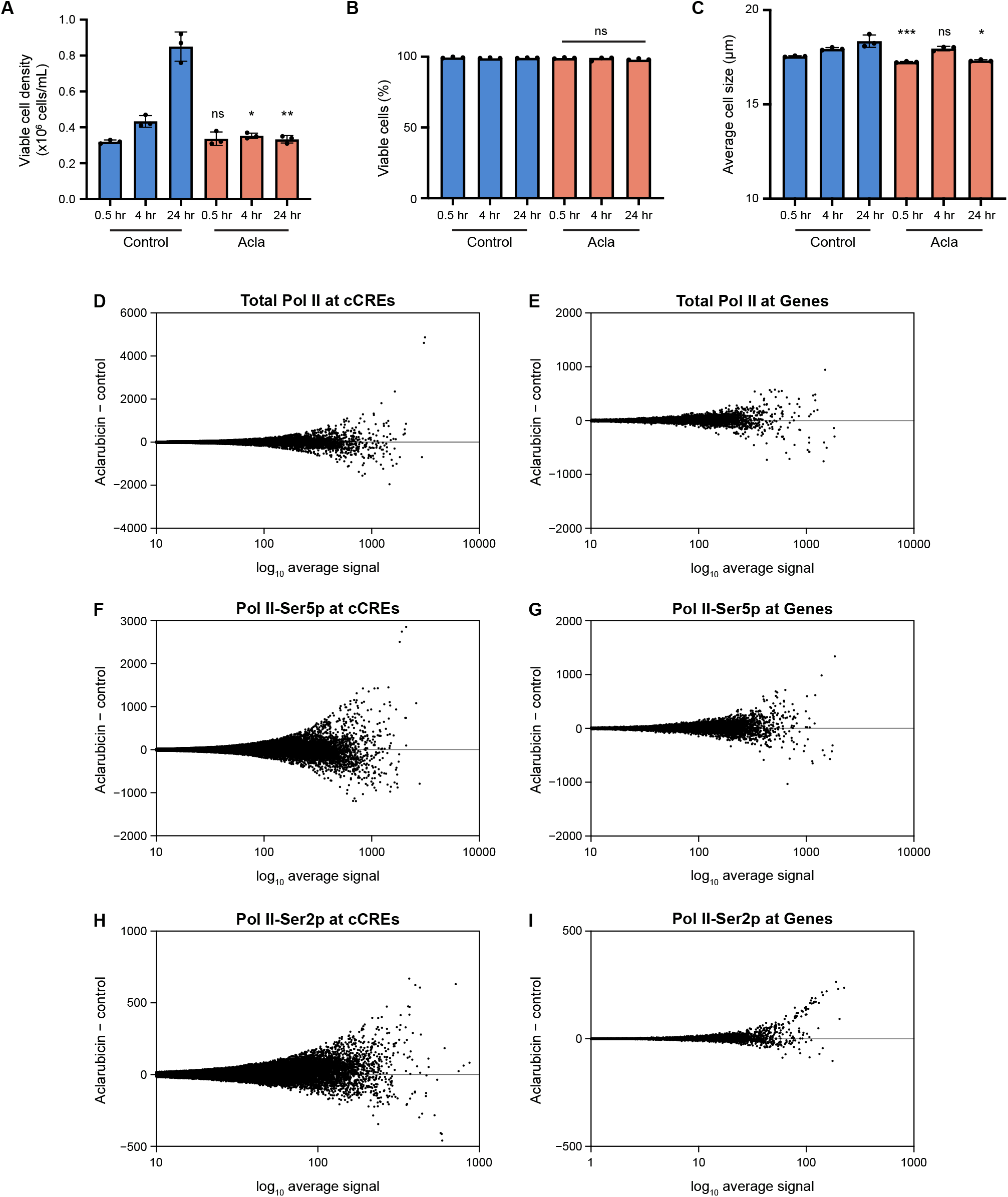
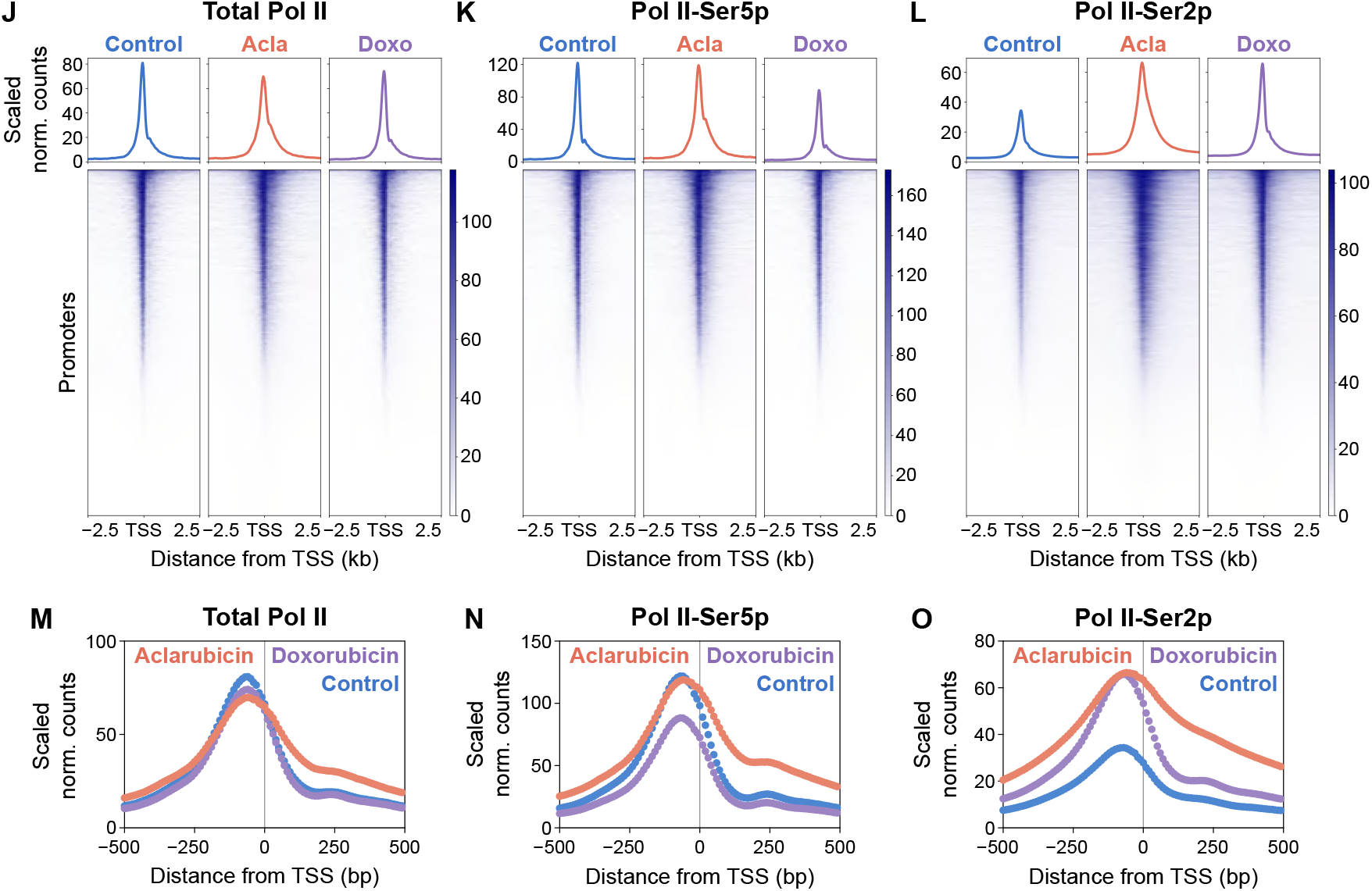
Impacts of drug treatment at various timepoints and changes in Pol II at cCREs and genes. **(A** to **C)** Viable cell density (A), viable cell population (B), and cell size (C) measurements after 30 minutes, 4 hours, and 24 hours treatment with vehicle control (DMSO) or aclarubicin (acla). For (A) to (C), bars represent mean ± SD of *N* = 3 biological replicates. Significant differences compared to control were calculated by unpaired twotailed Welch’s *t*-test. ns = not significant, \**P* < 0.05, \*\**P* < 0.01, *** *P* <0.001. **(D** and **E)** Aclarubicin-treated (*A*) minus control (*C*) total Pol II (Rpb1) normalized counts versus log_10_ (*A* + *C*)/2 over 2,343,081 annotated human cCREs (D) and 19,305 genes (E). **(F** and **G)** Aclarubicin-treated (*A*) minus control (*C*) Pol II-Ser5p normalized counts versus log_10_ (*A* + *C*)/2 over cCREs (F) and genes (G). **(H** and **I)** Aclarubicin-treated (*A*) minus control (*C*) Pol II-Ser2p normalized counts versus log_10_ (*A* + *C*)/2 over cCREs (H) and genes (I). **(J** to **L)** Average signal distribution plot and heatmap of total Pol II (Rpb1) (J), paused Pol II (Ser5p) (K), and elongating Pol II (Ser2p) (L) CUT&Tag signal under control, aclarubicin, and doxorubicin (doxo) treatment conditions aligned to the TSS of all promoters. For (J) and (K), merged data from two biological replicates per condition (control, aclarubicin-treated, and doxorubicin-treated) shown. For (L), merged data from eight biological replicates for control and aclarubicin-treated conditions and six biological replicates for the doxorubicin-treated condition shown. **(M** to **O)** Overlaid average signal distribution plot of total Pol II (Rpb1) (M), paused Pol II (Ser5p) (N), and elongating Pol II (Ser2p) (O) CUT&Tag signal under control, aclarubicin, and doxorubicin treatment conditions aligned to the TSS of all promoters.

**Fig. S2.**
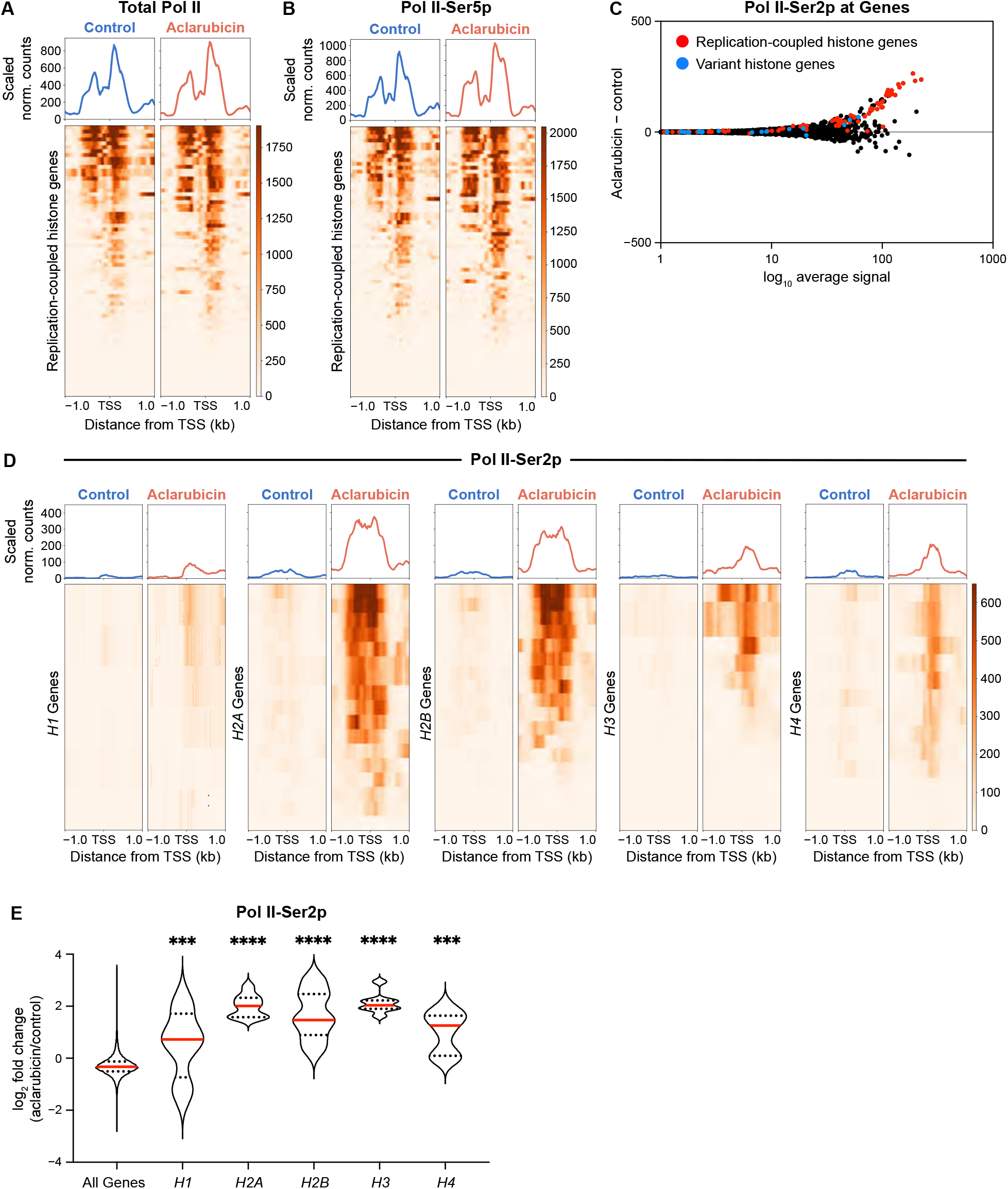
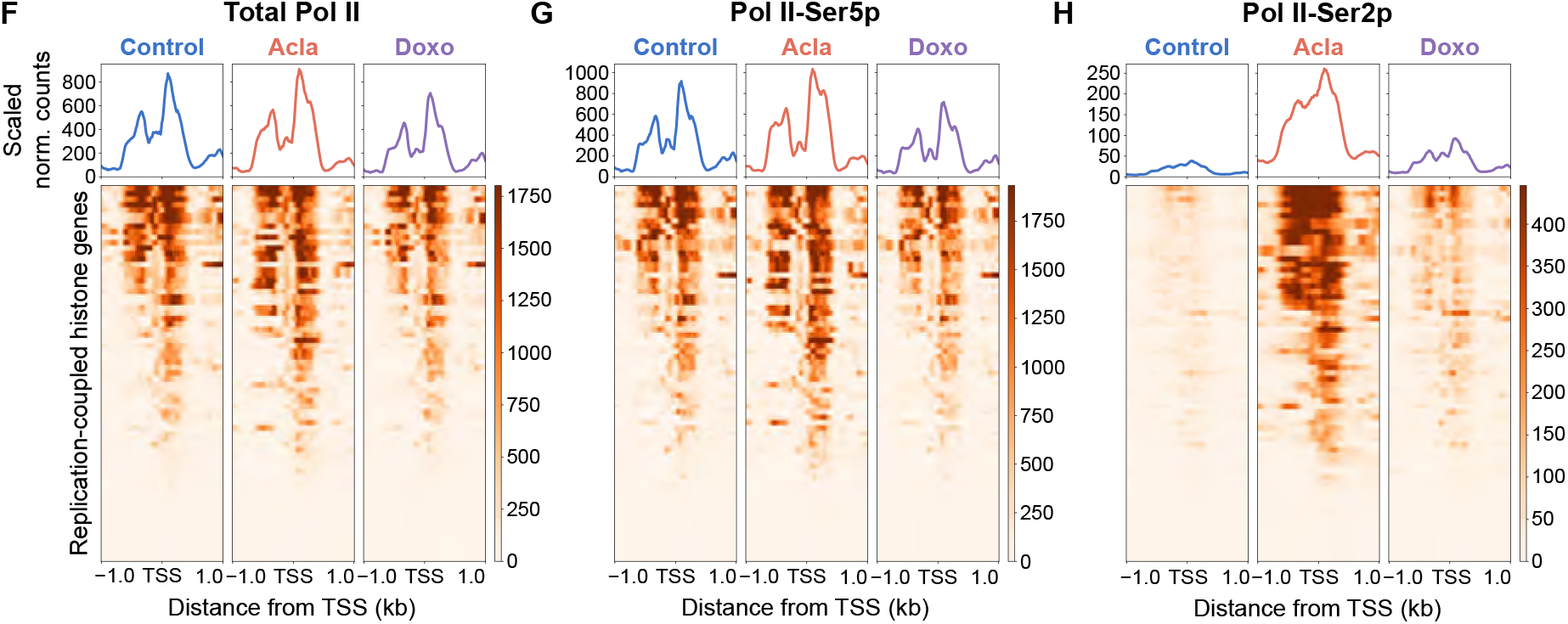
Impact of drug treatment on Pol II at the histone genes. **(A** and **B)** Average signal distribution plot and heatmap of total Pol II (Rpb1) (A) and paused Pol II (Ser5p) (B) CUT&Tag signal under control and aclarubicin treatment conditions aligned to the TSS of histone genes. (**C**) Aclarubicin-treated (*A*) minus control (*C*) Pol II-Ser2p normalized counts versus log_10_ (*A* + *C*)/2 across annotated human genes. Superimposed red dots indicate replication-coupled histone genes. Superimposed blue dots indicate variant histone genes. (**D**) Average signal distribution plot and heatmap of elongating Pol II (Ser2p) CUT&Tag signal under control and aclarubicin treatment conditions aligned to the TSS of individual histone genes. (**E**) Violin plots of per-gene log_2_ fold change (aclarubicin vs. control) of Pol II-Ser2p normalized counts across all genes and each histone gene class. Repeated-measures one-way ANOVA with Geisser-Greenhouse correction and Tukey’s multiple comparisons test was performed on per-replicate median log_2_ fold changes of drug–control pairs (*N* = 8). \*\*\**P* < 0.001, \*\*\*\**P* < 0.0001 versus all genes. Violin plots indicate median, minimum, maximum, and first and third quartiles. (**F** to **H**) Average signal distribution plot and heatmap of total Pol II (Rpb1) (F), paused Pol II (Ser5p) (G), and elongating Pol II (Ser2p) (H) CUT&Tag signal under control, aclarubicin, and doxorubicin treatment conditions aligned to the TSS of histone genes.

**Fig. S3.**
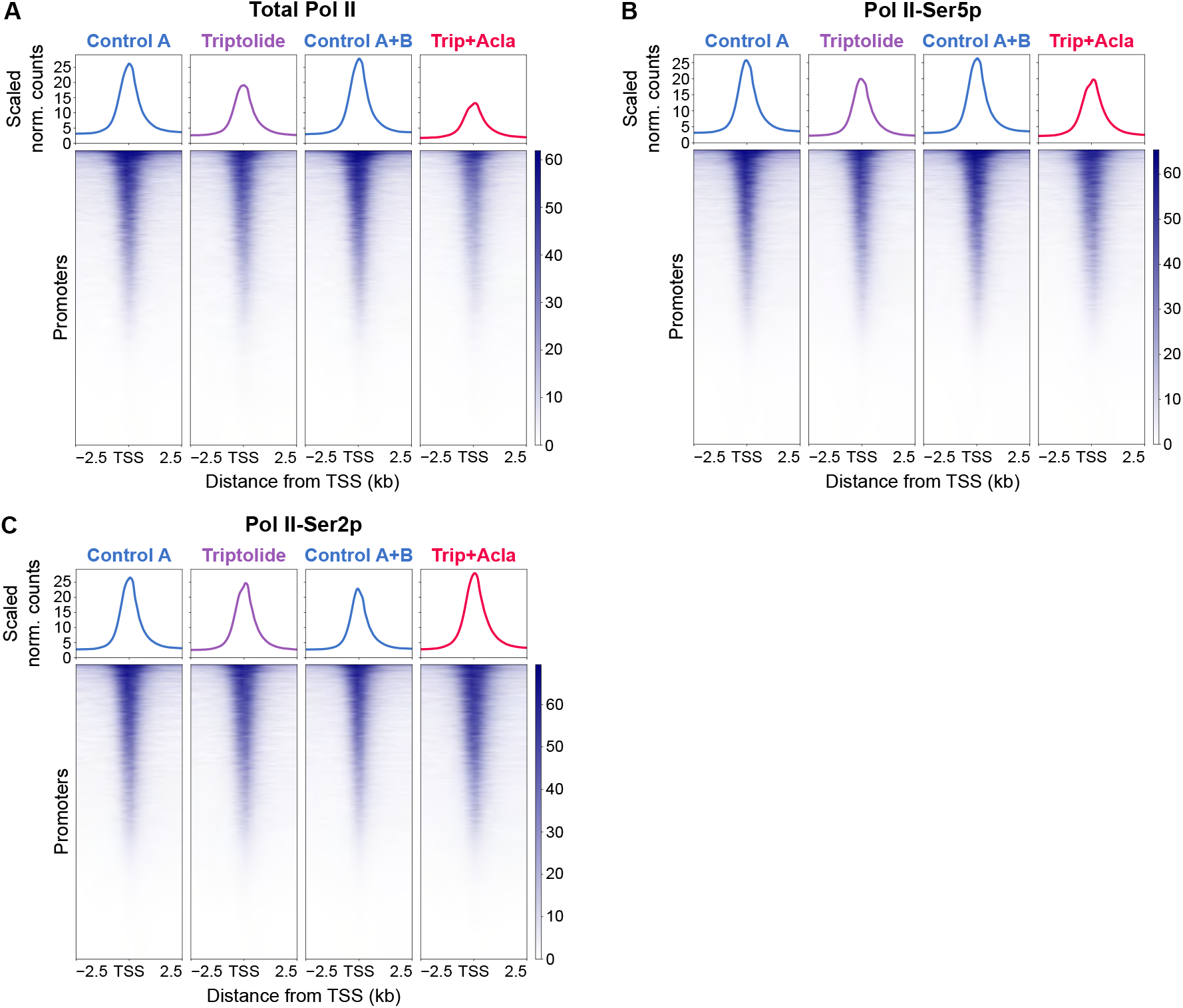
Pol II response to triptolide alone and in combination with aclarubicin. (**A** to **C**) Average signal distribution plot and heatmap of total Pol II (Rpb1) (A), paused Pol II (Ser5p) (B), and elongating Pol II (Ser2p) (C) CUT&Tag signal under treatment conditions aligned to the TSS of all promoters. For (A) to (C), merged data from three biological replicates per treatment condition shown.

**Fig. S4.**
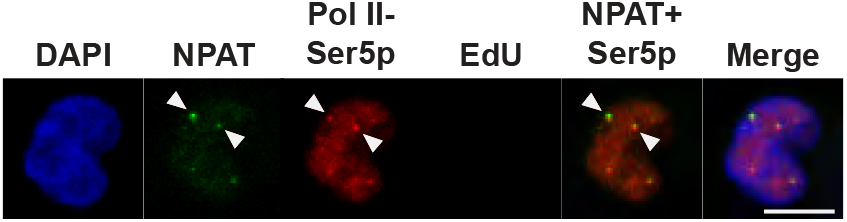
Colocalization of NPAT and Pol II-Ser5p foci in gap-phase cell. Representative immunofluorescence images of gap-phase K562 cell treated with vehicle control, stained for DAPI (blue), NPAT (green), Pol II-Ser5p (red), and EdU (magenta). Arrowheads indicate colocalizing NPAT and Pol II-Ser5p foci. Scale bar, 5 µm.

